# Human mitochondrial Lon protease initiates unidirectional degradation from either substrate terminus

**DOI:** 10.64898/2026.05.18.725877

**Authors:** Niko Schenck, Pavel Filipcik, Jan Pieter Abrahams

## Abstract

The ATP-dependent Lon protease (LonP1) unfolds and progressively degrades damaged or redundant proteins in the mitochondrial matrix. However, how LonP1 recognizes substrates remains unclear. Here, we investigated LonP1 degradation of the mitochondrial transcription factor TFAM, a physiological substrate. Using engineered TFAM variants carrying bulky moieties at either terminus, we show that degradation can initiate from either the N- or the C-terminus, with no evidence for internal initiation. Once engaged, substrate processing proceeds unidirectionally. Nanogold labeling experiments trapped TFAM termini within the N-domain cavity of LonP1, providing direct evidence that substrates enter through the N-domain assembly. A high-resolution cryo-EM structure of LonP1 in a transition-state-like complex with Mg^2+^, ADP, and AlF_3_ revealed substrate density within the axial channel with resolved main-chain carbonyl and Cα moieties, yet the peptide can be modelled equally well in either direction. Mass spectrometry of degradation products further supports initiation from either substrate terminus while confirming unidirectional processive translocation. These observations explain how LonP1 combines broad substrate specificity with selective recognition of flexible termini, avoiding initiation at internal flexible loops while enabling efficient ATP-dependent unfolding and degradation.

## Introduction

Mitochondria are essential organelles that sustain cellular energy production and coordinate diverse metabolic and signalling processes. To preserve organellar and cellular integrity, mitochondria rely on efficient protein quality control systems to remove damaged, misfolded, or redundant proteins. A central component of this system is the ATP-dependent Lon protease (LonP1), a conserved AAA+ (ATPases associated with diverse cellular activities) protease located in the mitochondrial matrix (Jadiya and Tomar, 2020; Rigas et al., 2009; Szczepanowska and Trifunovic, 2022).

As a member of the HCLR clade of protein unfoldases, LonP1 primarily degrades damaged proteins, particularly under oxidative stress (He et al., 2018; Ngo and Davies, 2009), but also regulates physiological substrates such as mitochondrial transcription factor A (TFAM) (Lu et al., 2013). As the primary architectural protein of mitochondrial DNA, TFAM dictates mitochondrial DNA copy number and transcriptional output. Consequently, precise control of TFAM levels by LonP1 is essential for mitochondrial biogenesis. Dysregulation of this proteostatic balance is a hallmark of ageing and human pathologies, including neurodegenerative disorders, cancer, and CODAS syndrome, a multisystem developmental disorder affecting skeletal and neurological health (Chan, 2020; Gibellini et al., 2018; López-Otín et al., 2013; Strauss et al., 2015).

Recent cryo-electron microscopy (cryo-EM) studies have revealed the functional structure of the LonP1 hexamer, defining a mechanism in which substrates are translocated through an axial channel via sequential, ATP-driven conformational changes. Substrate recognition has been proposed to involve the N-terminal domain, which forms a central cavity that connects to the axial channel responsible for substrate translocation, but direct experimental evidence for this model has been lacking (Mohammed et al., 2022; Shin et al., 2021). Despite these structural insights, the principles governing substrate recognition and the initiation of degradation remain unclear.

Unlike other cytosolic ATP-dependent proteases or homologues that rely on specific degradation tags, LonP1 appears to lack a universal degron motif. Instead, it operates within the protein-dense, oxidative environment of the mitochondrial matrix (Szczepanowska and Trifunovic, 2022; Taouktsi et al., 2022), where it must identify context-dependent features such as transiently exposed hydrophobic regions (Ondrovičová et al., 2005; Venkatesh et al., 2012). Furthermore, the human LonP1 enzyme features specialized ATPase arrangements and autoinhibitory elements that suggest a level of mechanistic complexity exceeding that of its bacterial homologs (Mohammed et al., 2022; Shin et al., 2021). This raises a fundamental question: how does a single protease recognize a wide range of structurally diverse substrates while maintaining the precision necessary for regulatory proteins such as TFAM? In addition, efficient ATP-dependent unfolding requires processive translocation of substrates through the LonP1 axial channel, implying that degradation must proceed directionally once initiated (Mohammed et al., 2022; Shin et al., 2021). How LonP1 reconciles this requirement with broad substrate recognition remains unknown. In particular, it is unclear whether substrate engagement is restricted to a specific terminus or whether LonP1 can initiate degradation independently of substrate polarity.

To address this question, we focused on the physiological LonP1 substrate TFAM and combined biochemical and structural approaches to probe substrate engagement and the initiation of degradation. We employed engineered TFAM variants with restricted terminal accessibility to define the requirements for substrate engagement. In parallel, we introduced N- and C-terminally Nanogold-labelled TFAM for single-particle cryo-EM localization of substrate interaction sites within the complex. Finally, to capture substrate engagement within the translocation machinery, we determined a cryo-EM structure of LonP1 stabilized in a transition-state-like conformation using Mg^2+^, ADP, and AlF_3_, an approach commonly used to trap ATP-dependent enzymes in defined functional states (Braig et al., 2000; Xu et al., 1997). Together, these approaches enable us to directly test how LonP1 engages substrates and initiates degradation at the molecular level.

### Resource availability

#### Lead contact

Jan Pieter Abrahams, Biozentrum, University of Basel, Basel, Switzerland & Paul Scherrer Institute, Laboratory of Multiscale Biology, Villigen, Switzerland.

#### Materials availability

Plasmids of LonP1 wild type and mutants of TFAM are available upon request. Otherwise, this study did not generate new unique reagents.

#### Experimental model and subject details

All recombinant protein constructs (human LonP1 and TFAM variants) were expressed in *E. coli* BL21 (DE3) cells. Cultures were grown in 2XYT medium at 37°C until induction.

#### Cloning and Mutagenesis

The human *LONP1* gene sequence encoding the mature protein (residues 125-959, lacking the mitochondrial targeting signal) was utilized for recombinant expression as previously described [27]. Plasmids for the physiological substrate TFAM (Human_TFAM_NoMTS_pET28) were obtained via Addgene (Addgene, product #34705). Custom pET-11a expression vectors harbouring TFAM terminal cysteine mutants were synthesized by GenScript Biotech Corporation. For the N-terminal mutant (TFAM_Nterm_Cys), the endogenous cysteine at position 221 was substituted with serine (C221S) to prevent internal labelling. This construct featured an N-terminal His-tag followed by a 3C protease cleavage site to facilitate tag removal. For the C-terminal mutant (TFAM_Cterm_Cys), the endogenous cysteine at position 7 was substituted with serine (C7S) and the construct retained an N-terminal His-tag for purification. These mutations ensured that each variant possessed only a single reactive cysteine at the desired terminus for functionalization with Monomaleimido-Nanogold® or PEG5K-Maleimide.

#### Protein expression and Purification

Recombinant human LonP1 and TFAM variants were expressed in *Escherichia coli* BL21 (DE3) cells. Cultures were grown in 2XYT medium at 37°C. LonP1 purification followed established protocols involving affinity and size-exclusion chromatography (SEC) (Mohammed et al., 2022). TFAM wild-type and mutant proteins were purified according to published procedures (Ngo et al., 2014) with modifications to preserve cysteine reactivity for downstream site-specific labelling. For TFAM mutants, reducing agents (e.g., DTT) were strictly omitted during SEC. The final SEC step was performed using a Superose 6 Increase 10/300 GL column equilibrated in HEPES buffer (50 mM HEPES, 150 mM NaCl, 5 mM MgCl_2_, pH 7.5). Purified proteins were supplemented with 10% (v/v) glycerol, flash-frozen in liquid nitrogen, and stored at -80°C.

#### In vitro protein degradation assay

Site-specific modification of TFAM variants was performed by incubating 5 µM single-cysteine mutants with 132 µM Methoxypolyethylene glycol maleimide 5 kDa (PEG5K-Maleimide; Sigma-Aldrich, product: 63187-1G) in reaction buffer (50 mM HEPES, 150 mM NaCl, 5 mM MgCl_2_, pH 7.5) for 30 min at room temperature. Labelling reactions were quenched with 3.3 mM 2-mercaptoethanol followed by a 20 min incubation at room temperature. Degradation was initiated by the addition of 0.45 µM LonP1 and 3 mM ATP to either labelled or unlabelled TFAM substrates. Aliquots were withdrawn at indicated time points and mixed 1:1 with SDS Gel Loading Buffer (0.13 M Tris-HCl, 0.2 M DTT, 15% glycerol, 0.02% bromophenol blue, 5% SDS, pH 6.8). All samples were denatured at 90°C for 5 min. Protein samples were analysed via SDS-PAGE using discontinuous NuPAGE™ 4-12% Bis-Tris Gel (Thermo Fisher Scientific, product: NP0322BOX) in NuPAGE™ MES SDS Running Buffer (Thermo Fisher Scientific, product: NP0002) at a constant 150 V for 60 min. Gels were visualized using EZBlue™ Gel Staining Reagent and compared against the PageRuler™ Plus Prestained Protein Ladder (Thermo Fisher Scientific, product: 26619).

#### Time-resolved mass spectrometry

Proteolysis reactions were initiated by incubating 13 µg LonP1 with either 8 µg TFAM or 8 µg casein (β-Casein from bovine milk; Sigma-Aldrich, product: C6905-250MG) in reaction buffer (50 mM HEPES, 150 mM NaCl, 5 mM MgCl_2_, 3 mM ATP, pH 7.5). Samples were withdrawn at the time points indicated. For sample preparation, a final concentration of 0.1% trifluoroacetic acid (TFA) in acetonitrile was added to the samples. The resulting peptide products were purified and concentrated using a C_18_ BioPureSPN MINI PROTO 300 columns (10-40 µL loading, 7-70 µg capacity; The Nest Group, Inc., product: HUM S18V) according to the manufacturer’s instructions for peptides. Before adding the samples, the columns were conditioned with acetonitrile and equilibrated twice using 0.1% TFA. The immobilised peptides were washed three times with 5% acetonitrile, 95% water (v/v), 0.1% TFA and eluted in two steps using 50% acetonitrile, 50% water (v/v), 0.1% TFA. The eluted peptides were dried and stored at -20 *°*C until further use.

#### LC-MS/MS analysis

Dried peptides were resuspended in 0.1% aqueous formic acid and subjected to LC-MS/MS analysis using an Orbitrap Fusion Lumos Mass Spectrometer fitted with an EASY-nLC 1200 (both Thermo Fisher Scientific) and a custom-made column heater set to 60 °C. Peptides were resolved using a RP-HPLC column (75 μm × 36 cm) packed in-house with C18 resin (ReproSil-Pur C18-AQ, 1.9 μm resin; Dr. Maisch GmbH) at a flow rate of 0.2 µL/min. The following gradient was used for peptide separation: from 5% B to 12% B over 5 min to 35% B over 40 min to 50% B over 15 min to 95% B over 2 min followed by 18 min at 95% B. Buffer A was 0.1% formic acid in water and buffer B was 80% acetonitrile, 0.1% formic acid in water. The mass spectrometer was operated in DDA mode with a cycle time of 3 s between master scans. Each master scan was acquired in the Orbitrap at a resolution of 120,000 FWHM (at 200 m/z) and a scan range from 375 to 1600 m/z followed by MS2 scans of the most intense precursors in the linear ion trap at “Rapid” scan rate with isolation width of the quadrupole set to 1.4 m/z. Maximum ion injection time was set to 50 ms (MS1) and 35 ms (MS2) with an AGC target set to 10^6^ and 10^4^, respectively. Only peptides with charge state 2-5 were included in the analysis. Monoisotopic precursor selection (MIPS) was set to Peptide, and the Intensity Threshold was set to 5^3^. Peptides were fragmented by HCD (Higher-energy collisional dissociation) with collision energy set to 35%, and one micro scan was acquired for each spectrum. The dynamic exclusion duration was set to 30 s.

#### Mass spectrometry data processing

For TFAM analysis, raw files were searched using the BioPharma Finder software suite v4.1 (Thermo Scientific) against the TFAMnoMTS sequence using the “Peptide Mapping Analysis” workflow. The default method for peptide mapping analysis was used with the following modifications: Mass Accuracy was set to 5 ppm, Minimum Confidence was set to 0.9 and Protease was set to Nonspecific. Data was exported on the component level, filtered for ID Type “MS2/Full” and MS1 intensities of all components belonging to the same peptide were summed up. For casein analysis, raw files were searched using MSFragger (v. 4.0) implemented in FragPipe (v. 21.1) against a *E. coli* database (consisting of 4390 protein sequences downloaded from Uniprot on 20220222) and 392 commonly observed contaminants spiked with the sequences of bovine casein (P02666) and human LonP1 (P36776) using the “Non-specific-peptidome” workflow with the following modifications: Variable modification of Cysteine was disabled, Peptide Prophet was enabled and MS1 quantification was enabled.

#### Site-specific Nanogold® labelling and cryo-EM of LonP1-TFAM complexes

##### Site-specific Nanogold® labelling

Single-cysteine TFAM mutants were site-specifically labelled with 1.4 nm Monomaleimido-Nanogold® (Nanoprobes, Inc., product: 2020A-5 × 6NMOL) for cryo-EM visualization. TFAM mutants were separately purified via size-exclusion chromatography (SEC) using a Superose 6 Increase 10/300 GL column to remove aggregates. Pure fractions of TFAM mutants (3 nmol) were incubated with 6 nmol Monomaleimido-Nanogold® in SEC buffer (50 mM HEPES, 150 mM NaCl, 5 mM MgCl_2_, pH 7.5) at a final concentration of 3 to 6 nmol/mL for 2 h at room temperature (according to the manufacturer’s protocol). Unbound gold particles were removed using a Superdex-75 column in SEC buffer. TFAM-Nanogold® conjugates were pooled and concentrated using a 10 kDa cut-off filter (Amicon® Ultra-0.5 mL Ultracel®).

#### Cryo-EM grid preparation

LonP1 was purified prior to sample preparation via size-exclusion chromatography (SEC) using a Superose 6 Increase 10/300 GL column to remove aggregates. For dataset I (C-terminal labelling), 0.8 mg/mL LonP1 were incubated with 1.5 µM TFAM_Cterm_Cys-Nanogold® and 3 mM ATP in SEC buffer for 45 min at room temperature. For dataset II (N-terminal labelling), 1.0 mg/mL LonP1 were incubated with 3.4 µM TFAM_Nterm_Cys-Nanogold® and 3 mM ATP in SEC buffer for 90 min at room temperature, since in vitro protein degradation showed reduced digestion rate for N-terminal labelled TFAM. Reaction mixtures (3.5 µL) were applied to glow-discharged Cu200 mesh R 0.6/1 Quantifoil Holey Carbon Grids. Dataset I grids were plunge-frozen in liquid ethane using a Vitrobot IV (Thermo Fisher Scientific) maintained at 10 °C (3 s blot time, 100% relative humidity, blot force -3). Dataset II grids were plunge-frozen using a Leica EM GP2 Automatic Plunge Freezer (Leica Microsystems GmbH) maintained at 10 °C (3 s blot time, 80% relative humidity, 75% nitrogen flow).

#### Cryo-EM data acquisition (Dataset I and II)

Dataset I and II were acquired on a Thermo Fisher Scientific Glacios TEM with a Gatan K3 direct Electron Detector operated at 200 kV. Movies were collected in dose-fractionated mode using SerialEM. For dataset I, a pixel size of 0.89 Å was used with a total electron exposure of 45.6⍰e^−^/Å^2^ over 40 frames (2.3 s exposure). For dataset II, the same pixel size was used with a total exposure of 48⍰e^−^/Å^2^ over 41 frames (2.3 s exposure). Real-time processing and data selection were performed in FOCUS (Biyani et al., 2017).

#### Image processing and 3D reconstruction

The subsequent processing was performed in CryoSPARC v4.1 (Punjani et al., 2017). For dataset I, particles were selected from motion-corrected and dose-weighted micrographs using the Blob picker with circular and elliptical blob. After two rounds of 2D classification, a subset of approximately 18’900 particles (from the initially 2.9 million picked particles) representing diverse orientations was used to train a Topaz 0.2.4 (Bepler et al., 2019) model for automated picking. The resulting 1.0 million particles underwent two rounds of 2D classification to identify 9’000 particles exhibiting both secondary structural features of LonP1 and Nanogold® density. These were utilized to train a second Topaz model specialized for Nanogold-containing particles. For Dataset II, Topaz was trained directly on Nanogold-containing particles identified after initial blob picking and 2D classification, resulting in a comparable particle selection. For both datasets, approximately 200’000 particles were extracted at 1.8 Å/pixel with a box size of 256 pixels after selection in one round of 2D classification. The extracted particles were subjected to an additional round of 2D classification using the ISAC (Iterative stable alignment and classification) algorithm implemented in the EMAN2/SPARX 1.2 software package (SPARX for High-Resolution Electron Microscopy) (Tang et al., 2007; Yang et al., 2012). Classification was limited to 300 images per class, a particle radius of 80 pixels and a 2.0 pixel error threshold. Particles clearly exhibiting Nanogold within the LonP1 N-domain cavity (15’000 for Dataset I and 10’000 for Dataset II) were selected, while those with multiple or non-specifically bound gold moieties were discarded. Selected particles were re-imported into CryoSPARC for ab-initio reconstruction and non-uniform refinement. To address structural heterogeneity in Dataset II, three-class ab-initio reconstruction was performed, and the largest class (approximately 7’000 particles) was selected for final refinement. In both datasets, focused refinement was performed using custom masks. Specifically, an N-terminal mask was used for particle subtraction of the gold-bearing N-domain, while a C-terminal mask was utilized for focused refinement of the protease core. Final coordinates from local refinements were used for 3D reconstruction of the complete protease-substrate complexes.

#### Atomic model fitting for Nanogold®-labelled complexes

Due to the high electron density of the 1.4 nm Nanogold® label relative to the LonP1 protein and its positional heterogeneity, particle alignment is impaired and the global resolution of the reconstructions was limited. Consequently, atomic modelling was restricted to rigid-body fitting of the protease core. The previously determined structure of the LonP1 P1a-state (PDB: 7NFY (Mohammed et al., 2022)) was utilized as a template. Rigid-body docking into the density maps was performed manually in Coot 0.9.8.96 (Emsley and Cowtan, 2004), followed by real-space refinement in Phenix 1.21 (Adams et al., 2010).

#### Analysis of non-specific Nanogold® binding

To evaluate potential non-specific interactions between the gold label and the protease, 0.36 nmol of purified LonP1 was incubated with 6 nmol non-functionalized Nanogold® (Nanoprobes, Inc., product: 2060-50NMOL) in SEC buffer (50 mM HEPES, 150 mM NaCl, 5 mM MgCl_2_, pH 7.5). Following 2 h incubation at room temperature, the mixture was subjected to size-exclusion chromatography using a Superose 6 Increase 10/300 GL column. Elution profiles were monitored simultaneously at 260 nm, 280 nm, and 420 nm to assess the migration of protein and gold particles.

#### Cryo-EM of Lonp1 trapped in a transition-state-like conformation

##### Preparation of ADP·AlF_3_-bound LonP1 complexes

To capture LonP1 in a transition-state-like conformation, LonP1 was subjected to size exclusion chromatography (SEC) using a Superose 6 Increase 10/300 GL column in order to remove aggregates. Pure fractions (1.0 mg/mL) were incubated with 1 mM ADP in SEC buffer (50 mM HEPES, 150 mM NaCl, 5 mM MgCl_2_, pH 7.5) for 20 min at room temperature. Subsequently, 3.125 mM NaF was added, followed by 0.625 mM AlCl_3_ after additional 20 min incubation for each step to generate the LonP1-ADP·AlF_3_ complex (Braig et al., 2000).

#### Cryo-EM grid preparation

The reaction mixture (3.5 µL) was applied onto glow discharged Cu300 mesh R 1.2/1.3 Quantifoil Holey Carbon Grids. Grids were plunge-frozen in liquid ethane using a Leica EM GP2 Automatic Plunge Freezer (Leica Microsystems GmbH) maintained at 10°C (3 s blot time, 80% relative humidity, 75% nitrogen flow).

#### Cryo-EM data acquisition (Dataset III)

Dataset III was collected on a Thermo Fisher Scientific Titan Krios G4 TEM operated at 300 kV, equipped with a Falcon 4i direct electron detector and a Selectris X TFS Imaging Filter. Movies were acquired using the EPU Software (Thermo Fisher Scientific) in dose-fractionated mode at a pixel size of 0.73 Å. A total electron dose of 40.5 e^−^/Å^2^ was applied over 52 frames (2.7 s exposure).

#### Image processing and 3D reconstruction

The subsequent processing was performed in CryoSPARC v4.1 (Punjani et al., 2017). Particles were selected from motion-corrected and dose-weighted micrographs using the Blob picker with circular and elliptical blob. An initial set of 2.6 million picks was subjected to three rounds of 2D classification. A representative subset of approximately 40’000 particles, selected for high resolution and diverse orientations, was used to train a Topaz 0.2.4 (Bepler et al., 2019) model for automated picking. The resulting 1.3 million particles underwent two rounds of 2D classification to identify around 750’000 particles for further refinement. To ensure map quality, the CryoSPARC Micrograph Junk Detector was applied to remove picks near contaminants, followed by orientation rebalancing, resulting in about 370’000 final particles. Ab-initio reconstruction was performed using eight classes, followed by heterogeneous refinement. This identified three distinct structural classes; two corresponding to the P1a-state conformation (180’000 particles) and one exhibiting a rotated N-domain characteristic of the P2a-state (77’000 particles). Each state was subjected to non-uniform refinement and further 3D classification to confirm conformational homogeneity. Local refinement was performed on the catalytic core using a mask for the ATPase/Protease domains.

#### Atomic model building and refinement for ADP·AlF_3_-bound LonP1

The atomic model of LonP1 bound to ADP·AlF_3_ was generated through iterative cycles of manual building in Coot 0.9.8.96 (Emsley and Cowtan, 2004) and automated refinement in Phenix 1.21 (Adams et al., 2010). The previously determined P1a-state (PDB: 7NFY (Mohammed et al., 2022)) served as initial templates. Prior to rigid-body fitting into the corresponding density map, ATPγS ligands were substituted with ADP·AlF_3_. Residues N-terminal to D412 were removed due to insufficient local resolution in the N-domain region. Refinement was performed using Phenix real-space refinement with a strategy consisted of five macrocycles including global minimization, local grid search, and individual isotropic ADP refinement. To maintain structural integrity custom bond restraints were implemented to stabilize the coordination between the P-loop backbone (residues 526-530) and the ADP·AlF_3_ ligands. The model included hydrogen atoms throughout the process to ensure optimal geometry and clash prevention. Due to the structural conservation and high degree of conformational similarity within the catalytic core across the identified 3D classes, the generated model was used as a template for the second density map, corresponding to a conformation similar to the P2a-state. After the same refinement strategy was applied, the N-domains from the P1a- and P2a-state were docked as rigid bodies into the corresponding densities using Coot 0.9.8.96 (Emsley and Cowtan, 2004) and Phenix 1.21 (Adams et al., 2010). Structural deviations between conformations were calculated using the superposition and RMSD tools in Coot and Phenix. Model quality was continuously monitored using MolProbity to address rotamer outliers and local fit-to-density issues. Final model statistics were generated using the comprehensive validation tools in Phenix.

## Results

### LonP1 initiates degradation at substrate termini

To determine whether LonP1 requires access to specific termini or can be initiated at internal sites, we generated single-cysteine mutants of the physiological LonP1 substrate TFAM and site-specifically modified these with a non-degradable polyethylene glycol (PEG5K) moiety at either the N- or C-terminus, or at both termini in the wild-type protein. Degradation of these substrates by LonP1 was monitored over time by SDS-PAGE and quantified by band intensity analysis (Figure 1 and S2).

**Figure 1:**
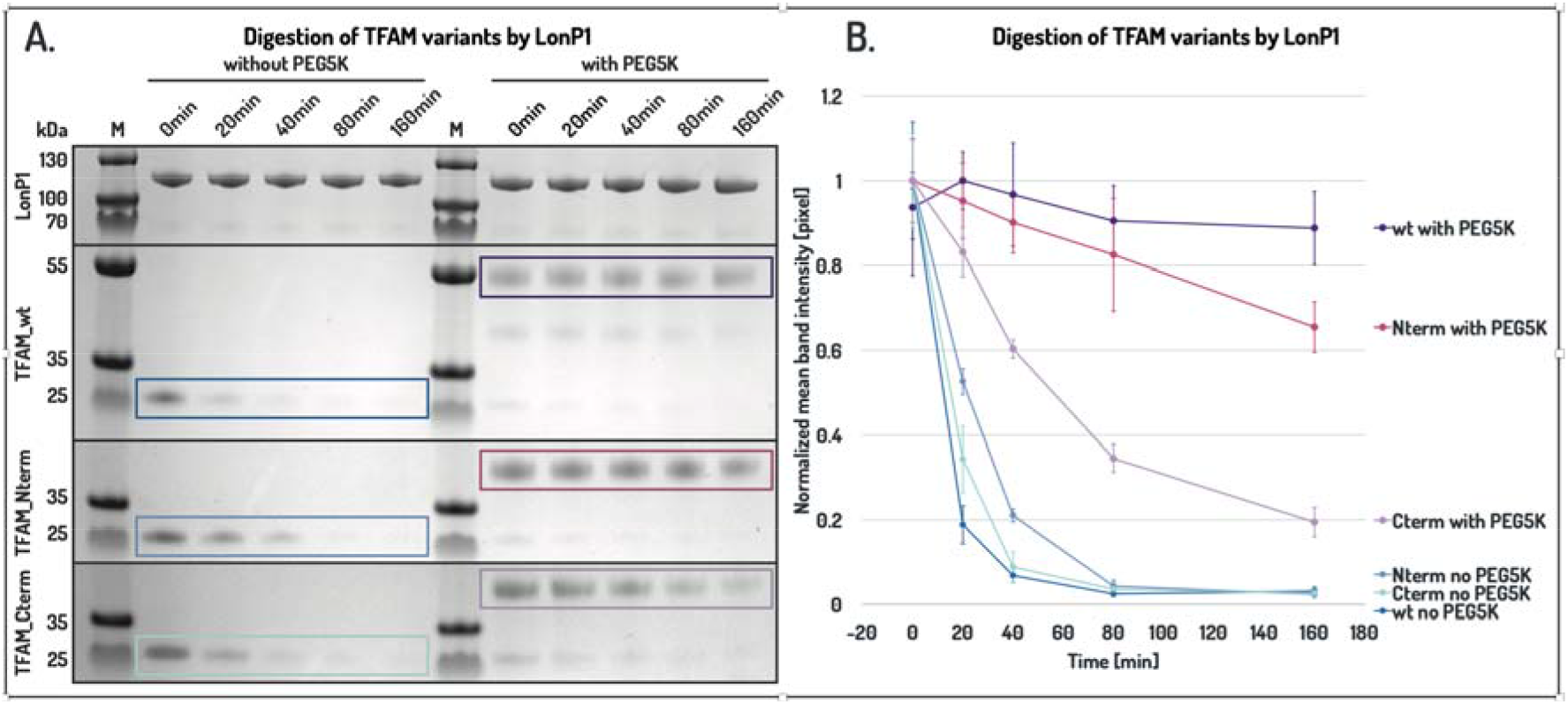
LonP1 preferentially initiates degradation from the N-terminus of TFAM. Time-dependent degradation of TFAM variants by LonP1 in the presence or absence of PEG5K modification at substrate termini. Nterm and Cterm denote N- or C-terminally modified single-cysteine mutants, respectively, whereas wt refers to the wild-type construct containing one cysteine at each terminus. **(A)** Representative SDS-PAGE analyses at indicated time points. Unlabelled TFAMs are outlined in shades of blue, and PEG5K-labelled TFAM variants in shades of purple. The full gel images are provided in Figure S2. **(B)** Quantification of band intensities from three independent experiments (mean ± standard error).

Unmodified TFAM was efficiently degraded by LonP1 in the presence of ATP, with the corresponding band disappearing within 20-40 min (Figure 1A). Similarly, both single-cysteine variants lacking PEG5K modification were degraded with comparable kinetics (Figure 1A, B), indicating that the introduced mutations do not impair substrate recognition. Modification of TFAM with PEG5K at a single terminus markedly affected degradation kinetics. N-terminally labelled TFAM exhibited strongly reduced degradation, with the corresponding band persisting throughout the 160 min time course and decreasing in intensity by about 40% (Figure 1B). In contrast, the unmodified counterpart was largely degraded within 20-40 min. C-terminal PEG5K modification also slowed degradation, but to a lesser extent, with band intensity decreasing to around 20% over the same period (Figure 1B). Strikingly, simultaneous modification of both termini in wild-type TFAM completely abolished degradation, as no significant decrease in band intensity was observed over 160 min (Figure 1A). These results demonstrate that LonP1 requires access to at least one free terminus for substrate engagement and that degradation is not initiated from internal regions of TFAM.

### Nanogold® labelling localises substrate entry

To determine where substrates enter LonP1, we employed site-specific Nanogold® labelling of TFAM variants carrying a single cysteine at either the N- or C-terminus. The bulky, electron-dense Nanogold® label enables selective identification of substrate-bound particles and, through steric trapping, allows localisation of substrate engagement within the complex. Representative 2D class averages showed a distinct high-density signal associated with the LonP1 hexamer, enabling selection of particle populations containing labelled substrate. Three-dimensional reconstructions of these particles revealed accumulation of the Nanogold® signal within the cavity of the N-domain at the entrance to the axial channel. Mapping the label onto the P1a-state of LonP1 positioned it directly above the translocation pathway, consistent with substrate engagement at the entry site (Figure 2).

**Figure 2:**
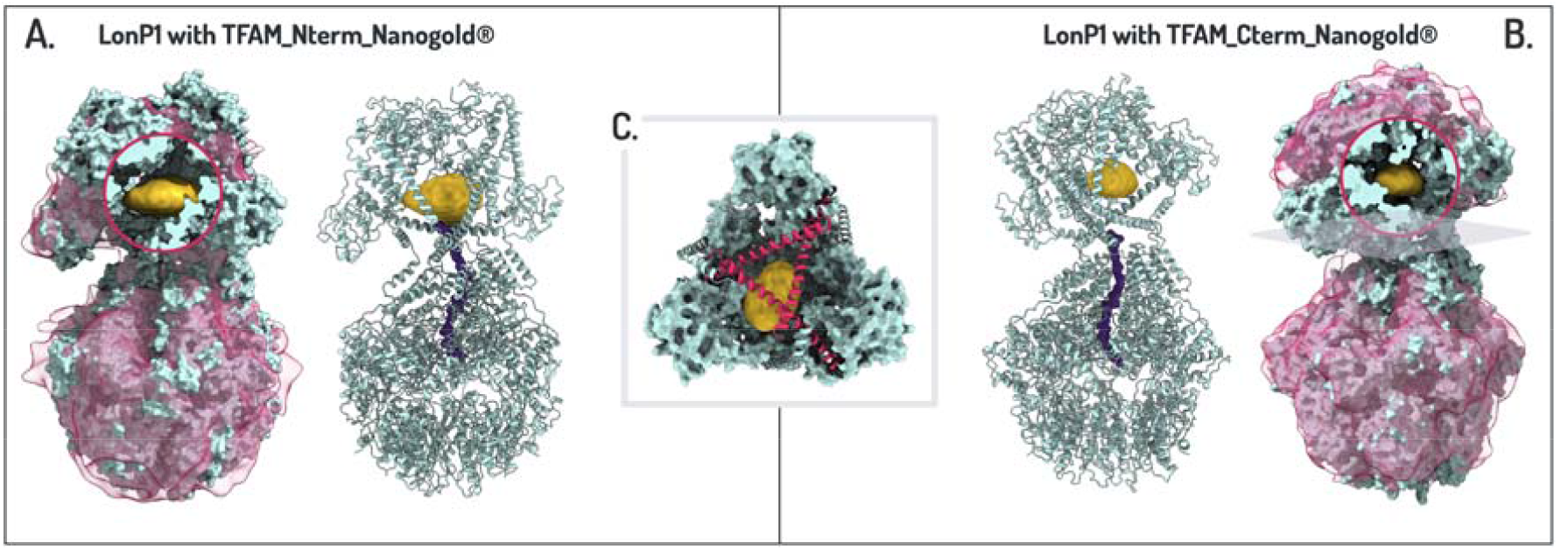
Nanogold® labelling localises substrate entry to the N-domain of LonP1. Cryo-EM analysis of LonP1 in complex with TFAM labelled with Nanogold® at either terminus reveals accumulation of the electron-dense label within the N-domain cavity at the entrance to the axial channel. **(A)** Density map of LonP1 bound to N-terminally labelled TFAM (pink), shown with rigid-body fit of the P1a-state (cyan). The Nanogold® density (gold) is positioned within the N-domain cavity (shown in the circled cross-sectional view). **(B)** Density map of LonP1 bound to C-terminally labelled TFAM, showing localisation of the Nanogold® signal at the same site. **(C)** View along the axial channel illustrating the position of the Nanogold® particle at the entrance to the translocation pathway.

Importantly, comparable localisation was observed for both N- and C-terminally labelled TFAM (Figure 2A, B), demonstrating that LonP1 can engage substrates from either terminus. In both cases, the Nanogold® density was positioned at the same site within the N-domain cavity, indicating that substrate entry occurs through a common structural pathway. The position of the Nanogold® signal along the axial axis is consistent with the experimental design, in which the bulky label is sterically trapped at the constriction marking the entrance to the translocation channel (Figure 2C).

Nanogold® labelling inherently limits achievable resolution, as the strong scattering signal of the gold particle dominates particle alignment and introduces positional heterogeneity. However, the purpose of this approach is not high-resolution structural reconstruction, but the direct localisation of substrate entry through steric trapping. In this context, the consistent positioning of the Nanogold® density across particle populations provides robust evidence that substrates are captured within the N-domain cavity prior to translocation. High-resolution structural insight into N- or C-terminal substrate engagement and translocation is provided by the transition-state-like LonP1-ADP·AlF_3_ complex described below.

Together with the biochemical data, these results support a model in which LonP1 engages accessible substrate termini and directs them into the axial translocation channel for processive degradation.

### LonP1 accommodates substrates in both orientations within the translocation channel

To obtain high-resolution structural insight into substrate engagement and translocation, LonP1 was stabilised in the presence of Mg^2+^, ADP, NaF and AlCl_3_ to generate ADP·AlF_3_, which mimics the γ-phosphate of ATP and traps NTP-hydrolysing enzymes in a transition-state-like configuration (Braig et al., 2000; Xu et al., 1997). Under these conditions, LonP1 adopts two closely related conformations resembling the previously described P1a and P2a processing states (Mohammed et al., 2022), differing by an approximately 60° rotation of the N-terminal domain assembly relative to the AAA^+^ domains (Figure S7A).

These datasets yielded a reconstruction at an overall resolution of 2.45 Å, enabling detailed analysis of both nucleotide occupancy and substrate density. ADP·AlF_3_ was clearly resolved at four inter-subunit catalytic sites, while the remaining two sites contained ADP (Figure S7C). Within the axial channel of the AAA^+^ domain, continuous poly-peptide density corresponding to a translocating substrate was observed spanning the translocation pathway (Figure S7B, D). This density exhibits regularly spaced features corresponding to C-alpha densities, but lacks resolvable side-chain density, consistent with a sequence-agnostic translocation channel. The quality of the density allowed modelling of a poly-alanine chain through the A-tunnel, that engages the YV-pincer motifs via backbone interactions. Notably, the density supports modelling of the substrate in both N-to-C and C-to-N orientations equally well. A poly-alanine chain can be fitted into the density in either direction, with Cα atoms aligning with the observed density features. In both orientations, backbone carbonyl groups form hydrogen bonds with the main-chain amides of the conserved residues Val566 and Tyr565 that constitute the YV-pincer. Real-space refinement yielded essentially indistinguishable statistics for both models (Figure 3).

**Figure 3:**
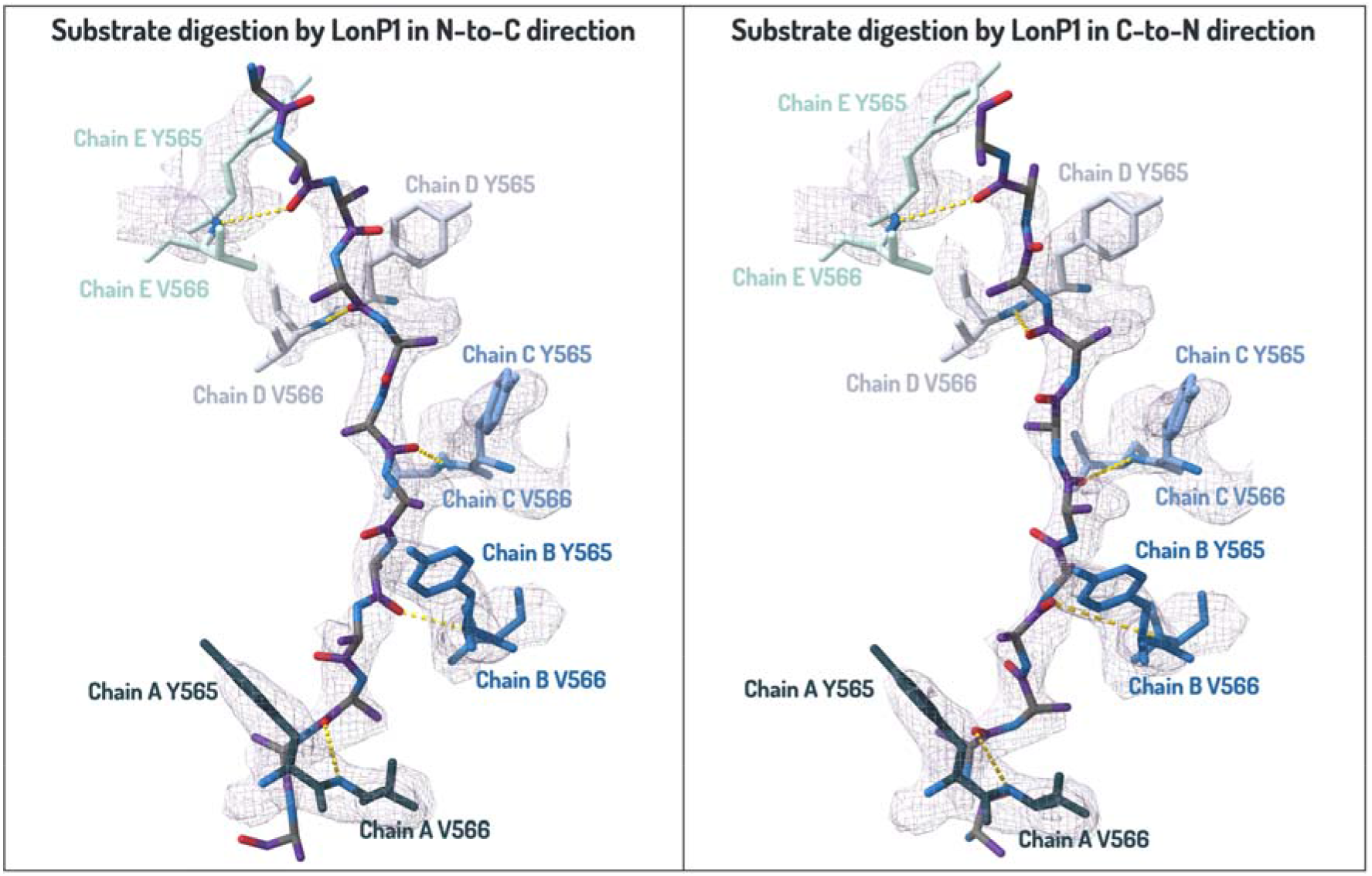
Bidirectional substrate engagement by the YV-pincer motifs in LonP1 stabilised with ADP·AlF_3_. The threaded polypeptide (purple) within the axial tunnel engages the YV-pincer region and can be modelled in both orientations. Residues of the respective subunits forming the pincer contacts are highlighted. The quality of the cryo-EM density enabled modelling of a poly-alanine chain along the substrate path, revealing backbone-mediated interactions with the conserved YV-pincer residues (yellow). The density is consistent with both N-to-C and C-to-N orientations, as a poly-alanine chain fits equally well in either direction, with close agreement between Cα positions and the cryo-EM map. In both models, backbone carbonyl groups are positioned to form hydrogen bonds with the main-chain amides between Val566 and Tyr565. Real-space refinement yielded indistinguishable statistics for both orientations, supporting bidirectional compatibility of substrate engagement.

These observations demonstrate that the LonP1 translocation machinery can accommodate polypeptide chains of either polarity. This structural plasticity provides a mechanistic basis for bidirectional substrate threading, consistent with the degradation behaviour observed for terminally modified substrates.

### Degradation directionality is substrate dependent

Time-resolved mass spectrometry was used to characterise the directionality of substrate degradation by LonP1. Peptide fragments generated during proteolysis of TFAM and casein were mapped onto their primary sequences (Figure 4).

**Figure 4:**
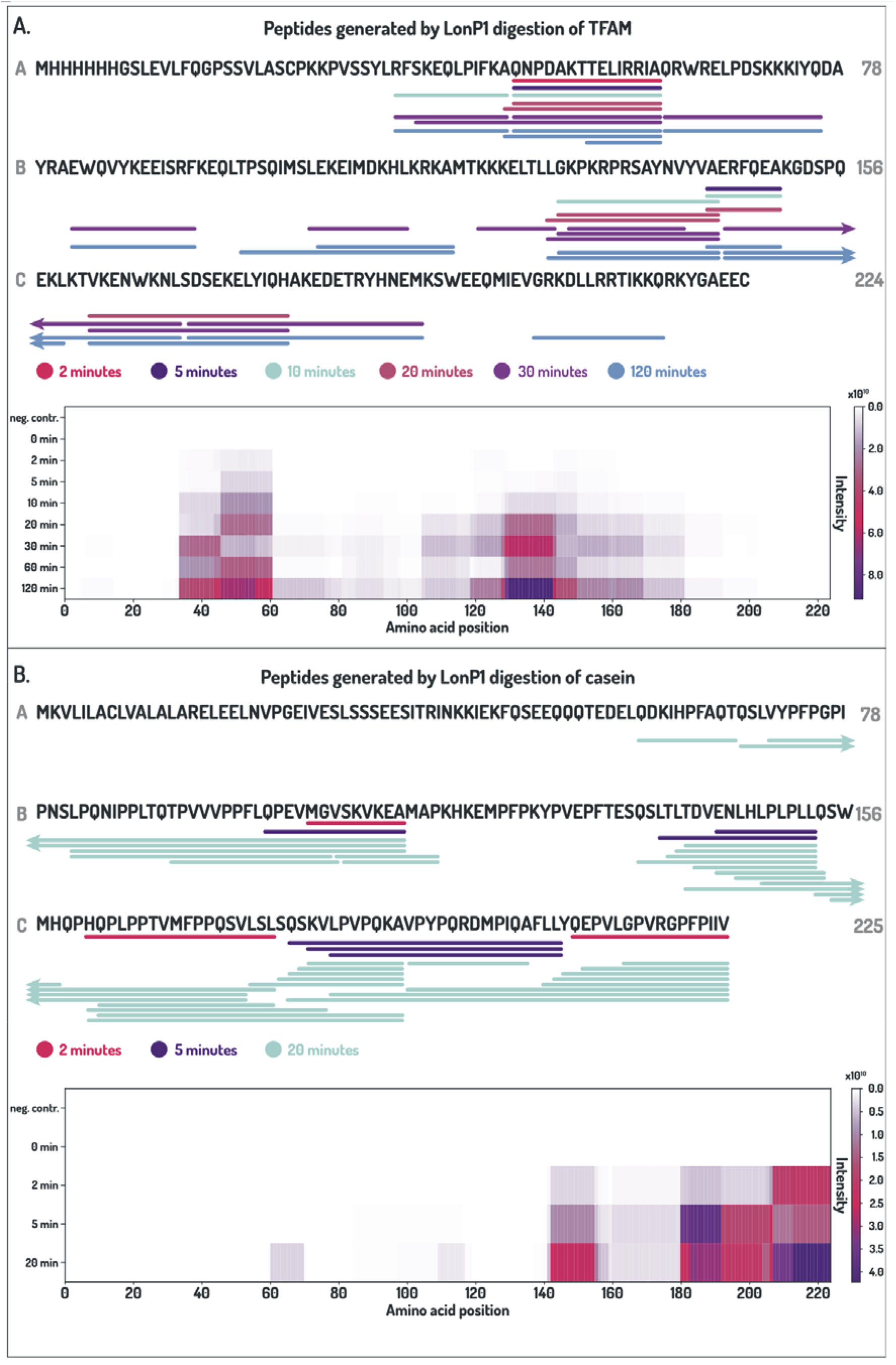
Time-resolved mapping of peptides generated during LonP1-mediated substrate degradation. Peptide fragments produced during degradation of TFAM and casein were identified by mass spectrometry and mapped onto their primary sequences. **(A)** TFAM. Peptides detected at the indicated time points are colour-coded by sampling time. Only peptides exceeding the intensity threshold (>10^7^) are shown (top). The heat map represents the cumulative intensity of all detected peptides (bottom). Control samples lacking LonP1 are shown for comparison. **(B)** Casein. As in (A), with peptides mapped onto the casein sequence.

For the physiological substrate TFAM, peptides originating from both terminal regions were detected at the earliest time points (2-5 min) (Figure 4A). The intensity of these terminal cleavage products increased over time, followed by the appearance of peptides from internal regions. However, peptides from central regions emerged largely simultaneously, precluding assignment of a defined direction of proteolytic progression. While terminally modified substrates could not be analysed by mass spectrometry due to interference of PEG, Nanogold® or fluorophore labels with peptide recovery and detection, the early appearance of cleavage products at both termini supports initiation from either end, followed by progressive fragmentation.

In contrast, degradation of the model substrate casein exhibited a more directional pattern (Figure 4B). Early cleavage products were predominantly derived from the C-terminal region, whereas peptides closer to the N-terminus appeared at later time points. This temporal progression indicates preferential initiation at the C-terminus, demonstrating that LonP1 can exhibit substrate-dependent directionality.

Analysis of peptide sequences from both substrates revealed a preference for cleavage after small hydrophobic residues, including alanine, valine and leucine, whereas charged residues were less frequently observed at cleavage sites. The diversity of newly generated N-termini and the occurrence of peptide ladders differing by only a few residues are consistent with processive degradation and limited sequence specificity.

## Discussion

Substrate degradation by LonP1 proceeds through a series of mechanistic steps, beginning with recognition, followed by engagement and threading into the translocation tunnel, and culminating in processive proteolysis.

Our data indicate that substrate recognition by LonP1 appears to be mediated by the N-domain assembly, which captures accessible terminal regions rather than internal sites. Notably, the presence of flexible internal segments is insufficient to support degradation, distinguishing LonP1 from AAA+ proteases capable of initiating unfolding at internal regions (Okuno et al., 2006). This behavior is consistent with steric constraints within the narrow LonP1 axial channel, which would preclude internal loop initiation and instead enforce terminal threading and processive unfolding (Mohammed et al., 2022; Shin et al., 2021). In this context, specificity arises from the ability to engage accessible polypeptide segments rather than from sequence-defined cleavage sites. Restricting initiation to terminal regions provides an efficient surveillance strategy, as these regions are often the first to become exposed upon partial unfolding, oxidative damage, or incomplete assembly (Jr and Trader, 2025; Parsell and Sauer, 1989; Varshavsky, 2011). This is consistent with the structural organisation of TFAM, which contains flexible N- and C-terminal segments. Together, these findings support a model in which the N-domain acts as an entry portal that selectively captures exposed termini, regardless of their polarity.

Following initial capture, the substrate must be threaded into the axial translocation channel. This step is constrained by the geometry of the pore and requires an unmodified polypeptide chain. The inability of PEG- and Nanogold-modified termini to support degradation indicates that bulky modifications prevent entry into the channel, consistent with a steric requirement for threading. The observation that even a flexible PEG moiety blocks degradation further suggests that threading depends on specific polypeptide backbone interactions within the pore. This is supported by the transition-state-like structure, which shows the substrate polypeptide engaging the conserved YV-pincer motifs through interactions with backbone carbonyl groups within the translocation tunnel. In contrast, the absence of resolved side-chain density indicates that side chains are accommodated within a relatively permissive pore environment, consistent with sequence-independent translocation.

Notably, reversal of substrate polarity changes the orientation of the peptide main-chain amides, but not the position of the peptide’s backbone carbonyl groups engaged by the YV-pincers. In the absence of additional orientation-specific interactions with the peptide’s main chain amides, the tunnel can therefore accommodate substrates in either N-to-C or C-to-N orientation without an intrinsic polarity preference.

Once engaged, substrate translocation and degradation proceed in a processive and unidirectional manner. The direction of degradation is therefore determined by the orientation of the substrate at the moment of engagement rather than by an inherent polarity preference of the translocation machinery. This distinction between flexible initiation and committed translocation reconciles the ability of LonP1 to process substrates from either terminus with the requirement for directional, ATP-driven unfolding and degradation. To our knowledge, this provides one of the clearest demonstrations that a AAA+ protease can support unidirectional processive translocation independently of substrate polarity.

The efficiency and direction of degradation are further influenced by substrate-specific properties. In the case of TFAM, accessible terminal regions allow initiation from either end, while more structured regions may slow or transiently impede translocation. In contrast, the preferential processing of casein from one terminus suggests that asymmetries in disorder or stability can bias the site of engagement (Figure S1E, F). Thus, substrate architecture can modulate degradation behaviour without altering the underlying mechanism (Lee et al., 2001).

Finally, mass spectrometry reveals a relatively permissive proteolytic specificity, with a preference for small hydrophobic residues and evidence of processive trimming. This is consistent with a model in which substrate recognition is governed by structural accessibility, while cleavage within the proteolytic chamber proceeds with limited sequence constraint (Lee et al., 2001; Ondrovičová et al., 2005; Venkatesh et al., 2012).

The ability of LonP1 to initiate degradation from either terminus has important implications for the regulation of physiological substrates such as TFAM. TFAM undergoes substantial conformational changes upon DNA binding, with terminal regions becoming less accessible in the assembled nucleoprotein complex (Figure S1A-D) (Ngo et al., 2014, 2011). Consistently, LonP1 targets TFAM in its unbound or partially unfolded state, while DNA-bound TFAM is protected from degradation (Lu et al., 2013).

More generally, the ability to engage both N- and C-termini allows LonP1 to recognise structurally diverse substrates via accessible, disordered terminal regions rather than sequence-specific motifs. Coupled to committed, processive degradation once engaged, this provides an efficient mechanism for protein quality control while avoiding indiscriminate cleavage of exposed polypeptide segments.

## Supplemental data

**Figure S1:**
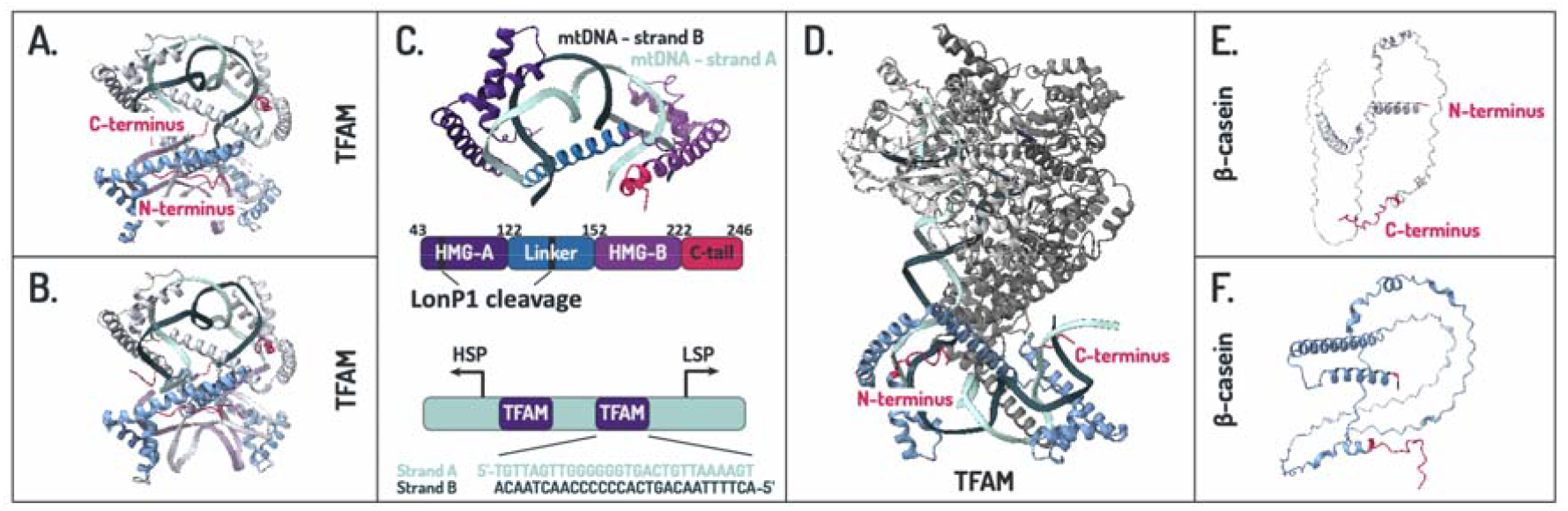
Structural context of TFAM and β-casein in relation to substrate accessibility for LonP1. Available structures of human TFAM in complex with mitochondrial DNA and structural models of bovine β-casein illustrate differences in terminal organisation that may influence substrate engagement by LonP1. Protein backbones are shown in shades of blue, with N- and C-termini highlighted in pink. Nucleic acids are presented in shades of lila or green, and additional transcription complex components are given in grey. **A and B**. Cryo-EM structures of TFAM bound to mitochondrial DNA (PDB: 4NOD and 4NNU) demonstrate compaction of TFAM upon interaction with the light strand promoter (Ngo et al., 2014). **C**. Structural overview of the TFAM-mtDNA complex with a schematic representation of the arrangement of mature TFAM in the same colour code below. At the bottom, the arrangement of the LSP and HSP1 promoters are shown. TFAM-binding sites are oriented oppositely relative to the transcription direction; the LSP fragment from the TFAM-mtDNA complex is indicated (adapted from (Ngo et al., 2011)). **D**. Structure of TFAM within the mitochondrial transcription initiation complex (PDB: 9MN4), illustrating tight wrapping of TFAM around DNA during transcription initiation (Herbine et al., 2025). **E and F**. AlphaFold (AF-P02666-F1) and AlphaFill (P02666) models of casein show a largely disordered polypeptide with limited helical content (Jumper et al., 2021). The N-terminus begins with a helical segment, whereas the C-terminus contains an extended flexible region.

**Figure S2:**
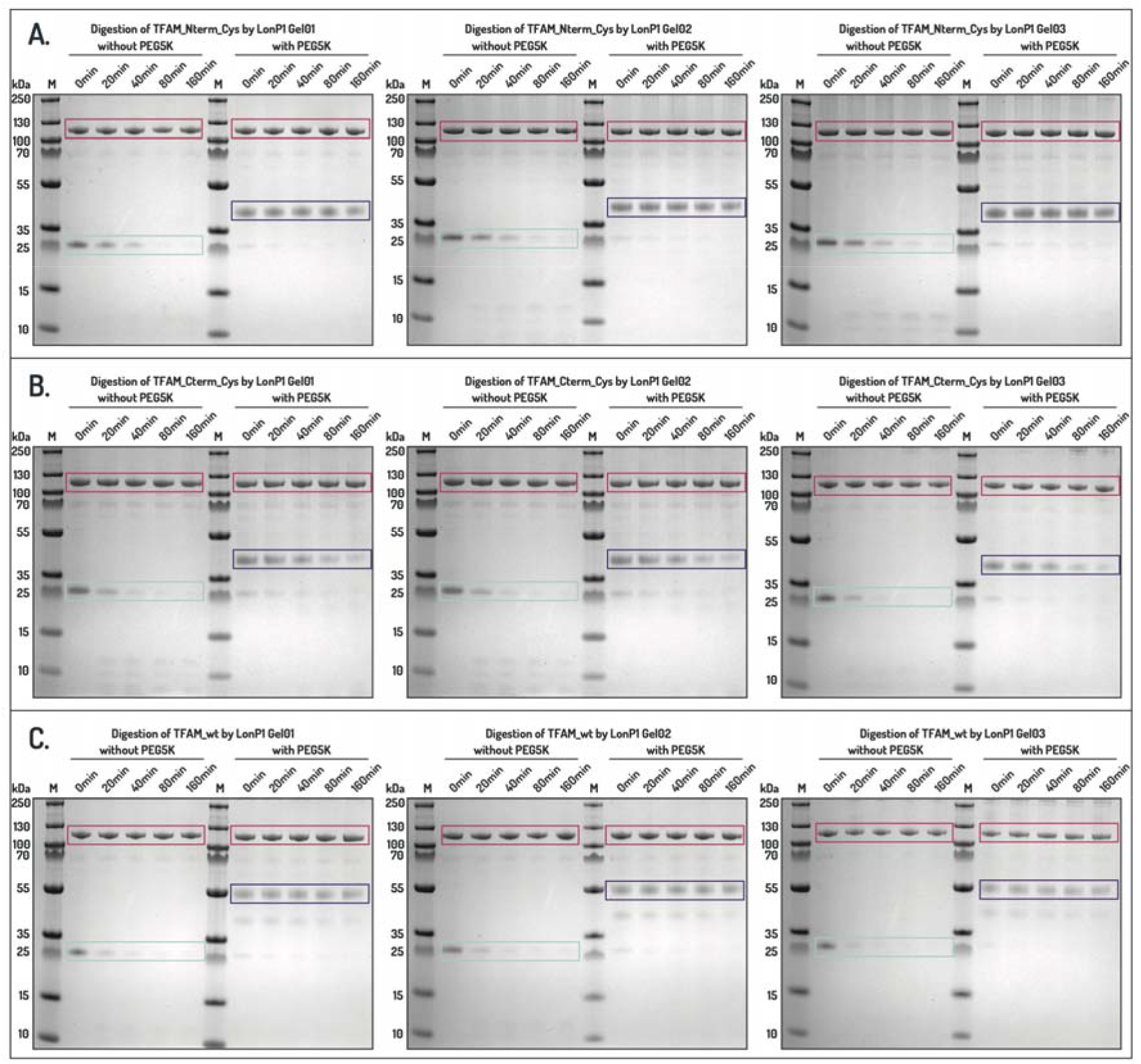
SDS-PAGE analysis of LonP1-mediated degradation of terminal labelled TFAM variants. The requirement for a free substrate terminus in LonP1-mediated degradation was assessed by monitoring the time-dependent digestion of TFAM variants in the presence or absence of a non-degradable polyethylene glycol (PEG5K) label. Samples were analysed by SDS-PAGE at the indicated time points. Bands corresponding to unlabelled TFAM are outlined in cyan, PEG5K-labelled TFAM species are outlined in purple, and LonP1 bands are indicated in pink. Experiments were performed in triplicate. **A**. TFAM variant containing a single cysteine at the N-terminus, analysed with and without PEG5K modification. **B**. TFAM variant containing a single cysteine at the C-terminus, analysed with and without PEG5K modification. **C**. Wild-type TFAM, containing one native cysteine residue at each terminus, analysed for degradation by LonP1 in the absence and presence of PEG5K modification at both termini.

**Figure S3:**
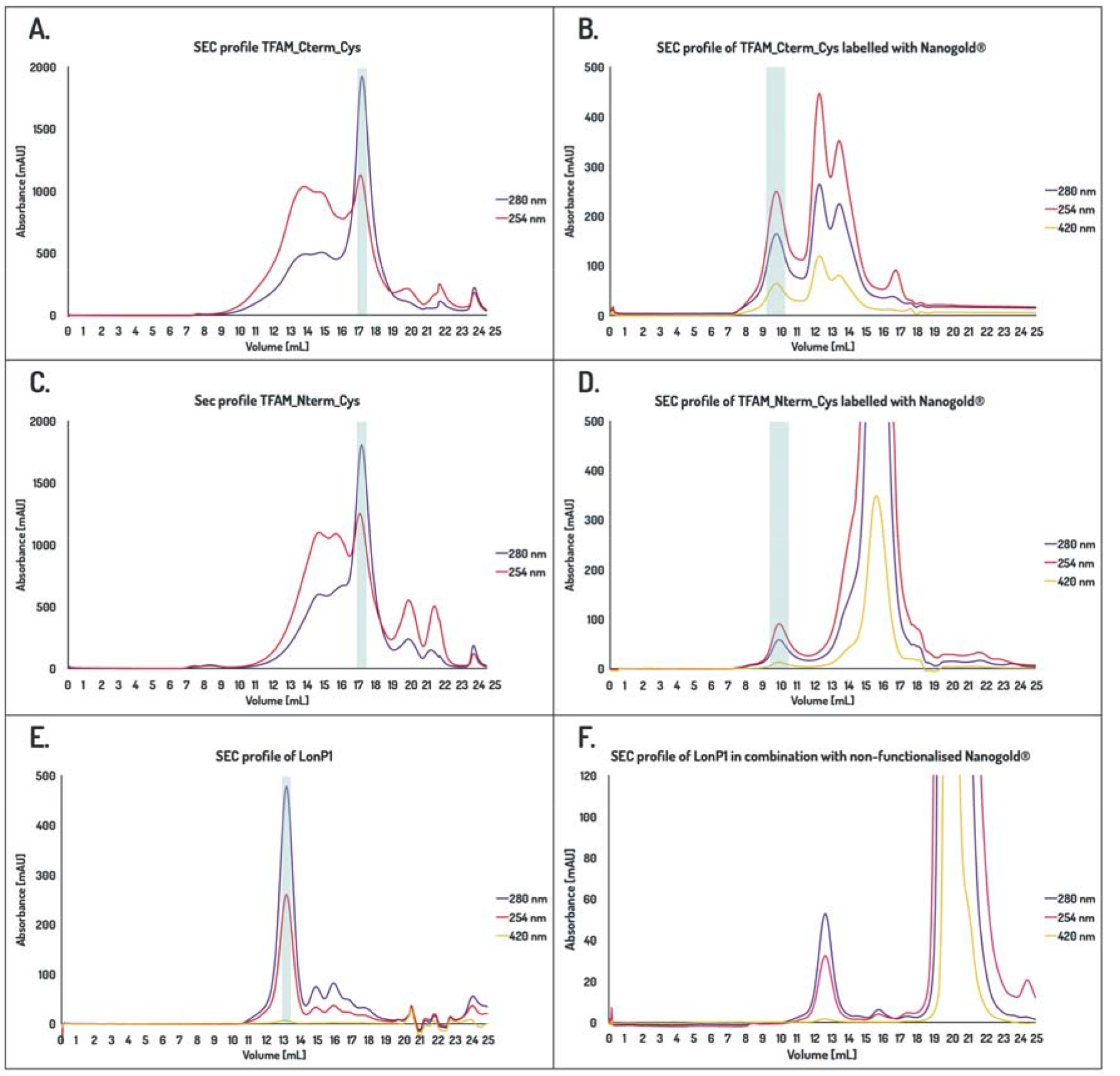
Cryo-EM sample preparation for Datasets I and II. As a final purification step samples were passed through a size exclusion column. For purification of target protein, a Superose 6 Increase 10/300 GL column was used, while unbound gold and TFAM-gold conjugates were separated with a Superdex-75 column. In both cases size exclusion buffer was used. (50 mM HEPES, 150 mM NaCl, 5 mM MgCl_2_). **A**. Size exclusion (SEC) profile of purified TFAM mutant featuring a single cysteine at the C-terminus. The fractions used are indicated in cyan. **B**. After labelling with Nanogold® the bound and unbound fractions were separated by size exclusion. **C**. Size exclusion profile of purified TFAM mutant featuring a single cysteine at the N-terminus. The fractions used are indicated in cyan. **D**. After labelling with Nanogold® the bound and unbound fractions were separated by size exclusion. **E**. Size exclusion profile of human wild type LonP1 used for cryo-EM sample preparation. **F**. In order to rule out unspecific binding of gold beads to LonP1, we incubated LonP1 with non-functionalized Nanogold® and analysed the sample using size exclusion chromatography.

**Figure S4:**
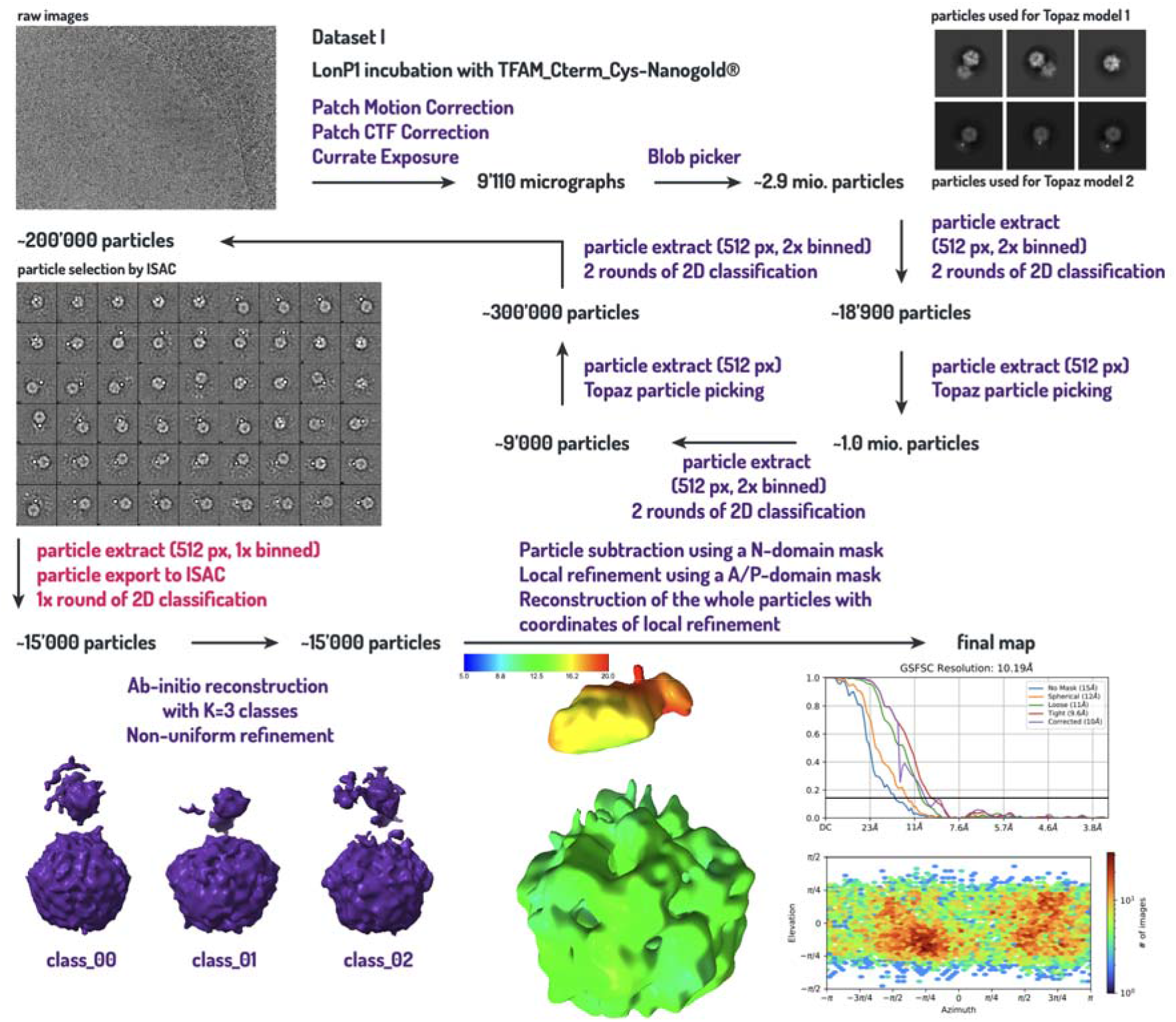
Data collection and 3D density calculation of LonP1 with TFAM_Cterm_Cys-Nanogold®. Summary of the results obtained from Dataset I, resulting in LonP1 with additional density of Nanogold®-labelled substrate protein inside its N-terminal cavity. After carefully selection of particles with a gold signal inside the N-domain, positioned at the entrance to the translocation tunnel, a total of 15’380 particles underwent the final refinement. The local resolution map together with the FSC and angular distribution plots of this map are given.

**Figure S5:**
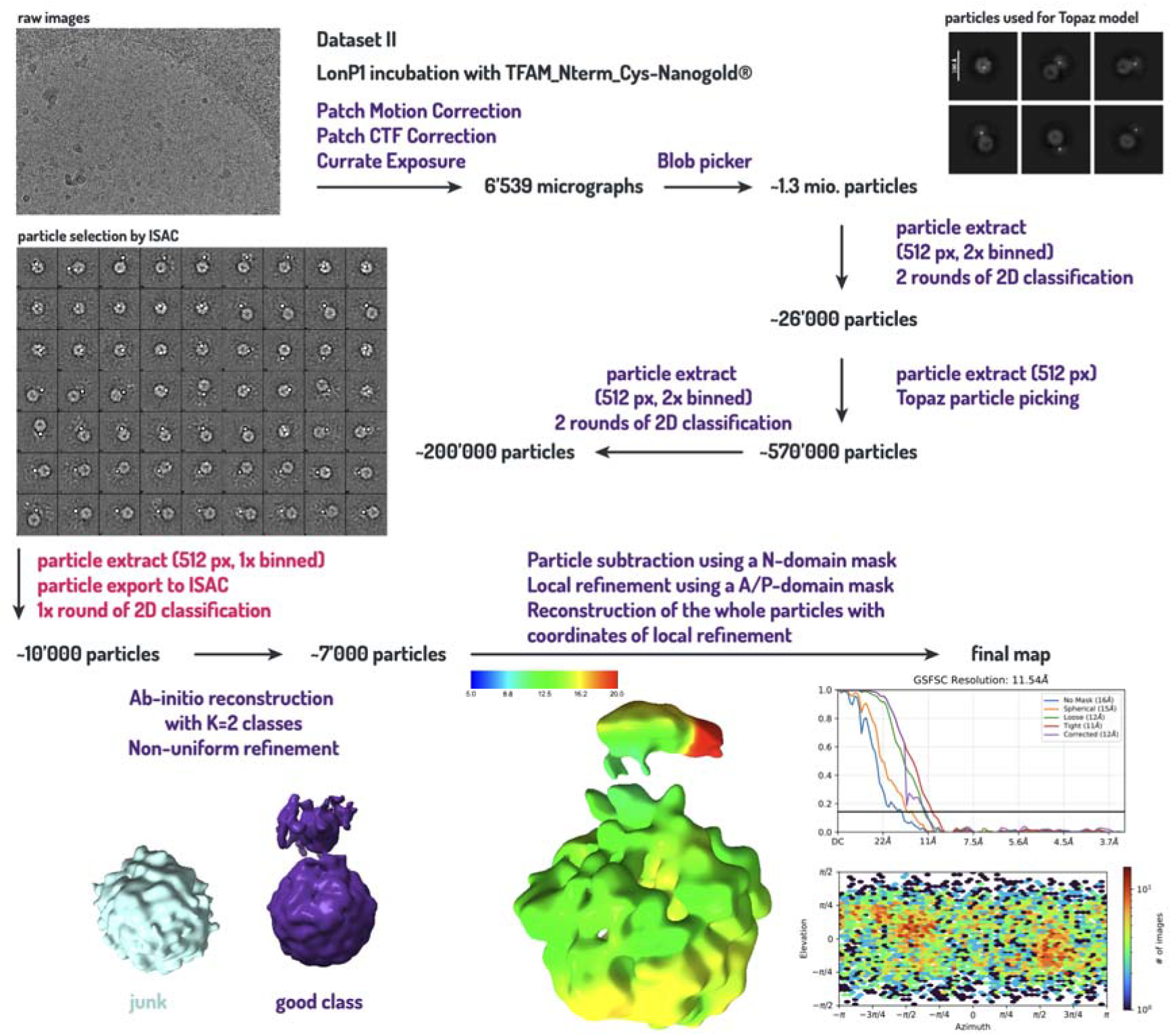
Data collection and 3D density calculation of LonP1 with TFAM_Nterm_Cys-Nanogold®. Summary of the results obtained from Dataset II, resulting in LonP1 with additional density of Nanogold®-labelled substrate protein inside its N-terminal cavity. After carefully selection of particles with a gold signal inside the N-domain, positioned at the entrance to the translocation tunnel, a total of 7’385 particles underwent the final refinement. The local resolution map together with the FSC and angular distribution plots of this map are given.

**Figure S6:**
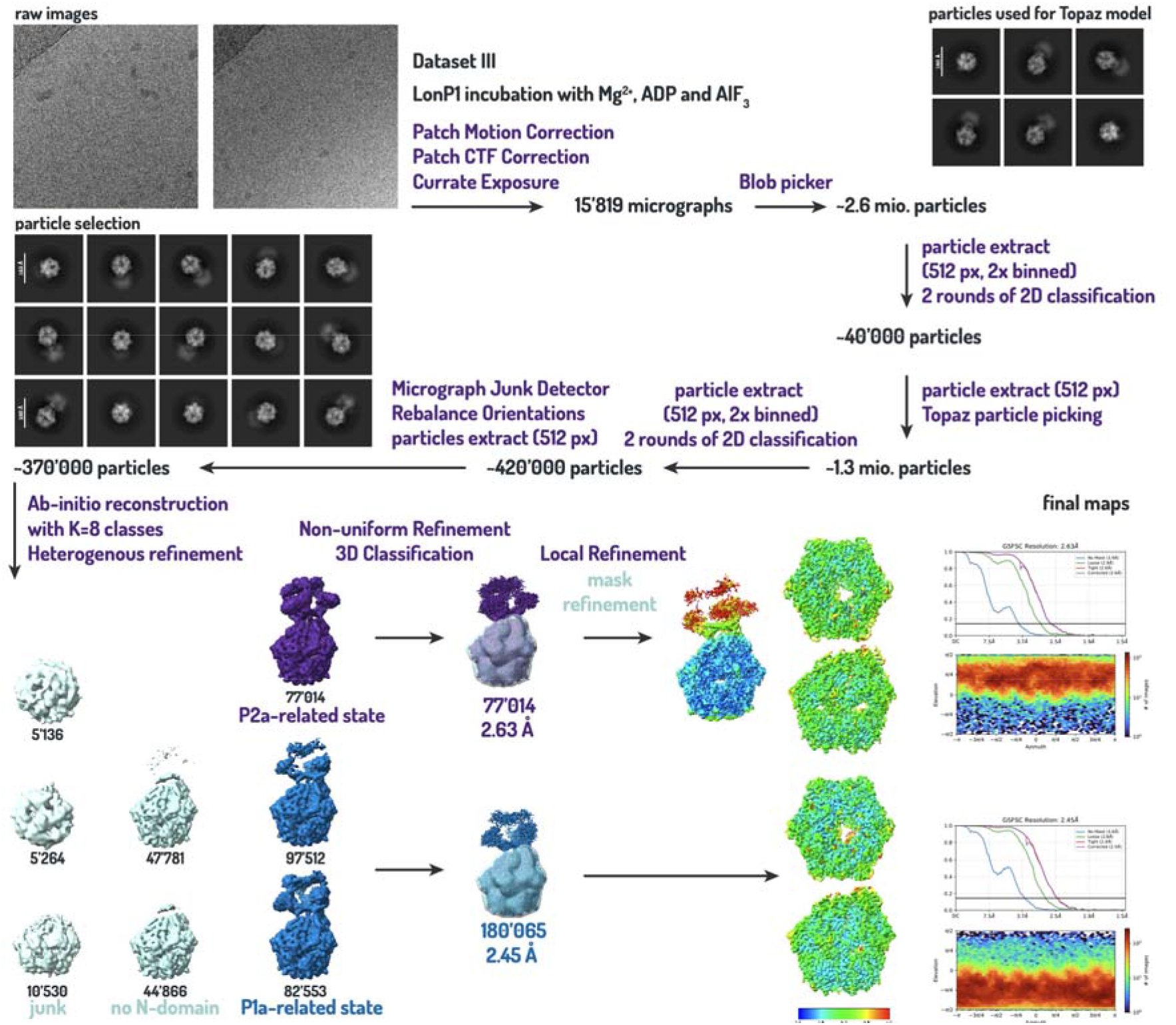
Data collection and 3D density calculation of LonP1 stabilised with ADP·AlF_3_. Summary of the results obtained from Dataset III, resulting in two conformations of LonP1 closely related to the P1a/P2a-states, but bound to ADP·AlF_3_ and local differences. After 3D classification the two distinct conformations could be identified by their respective orientation of the N-domain towards the proteolytic core. The map resolutions and the quality were further improved (The A- and protease soft mask is shown in cyan). The presence of Mg^2+^, ADP and AlF_3_ improved structural ordering and the final resolution of the maps (P1a-related state: 2.45 Å; P1a-related state: 2.63 Å). The local resolution map together with the FSC and angular distribution plots of this map are given.

**Figure S7:**
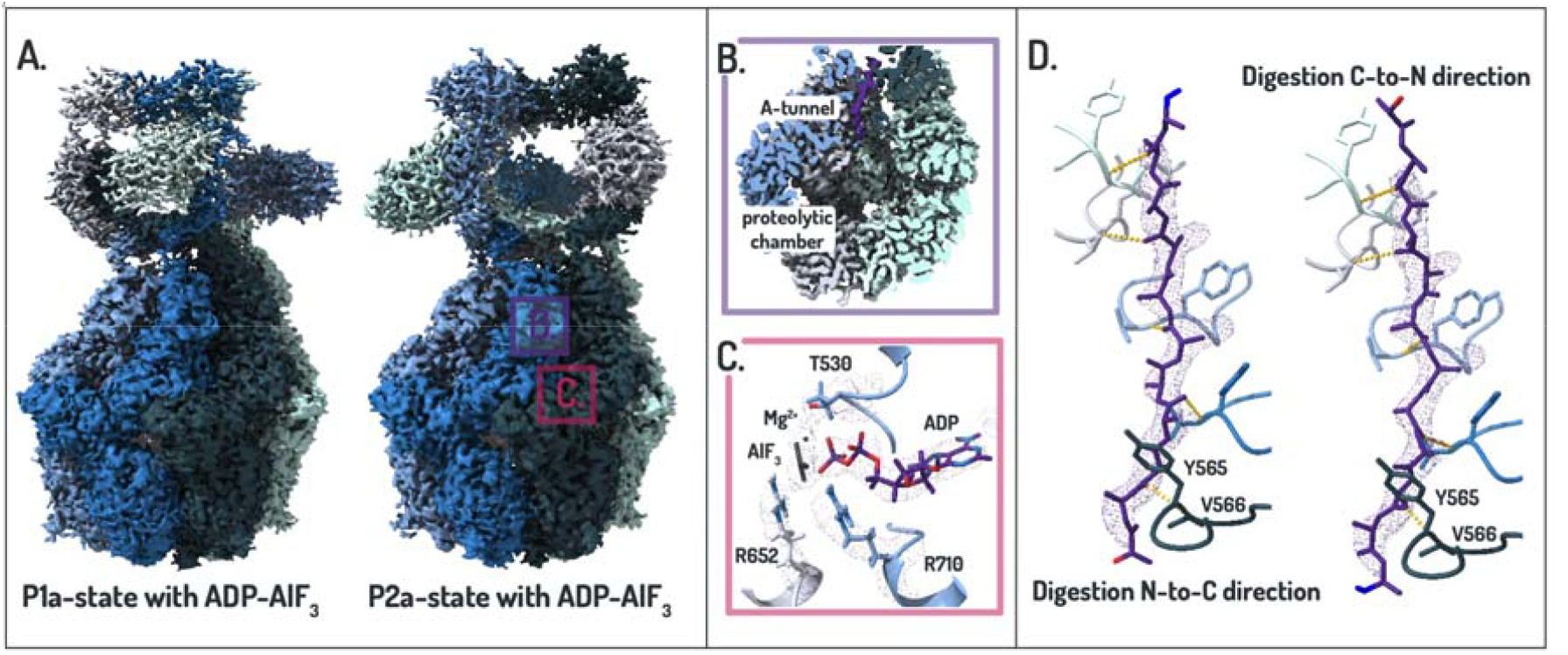
High-resolution structure of LonP1 stabilised with ADP·AlF_3_ reveals substrate density in the axial channel. Cryo-EM structure of LonP1 in the presence of Mg^2+^, ADP and AlF_3_. **(A)** Overall architecture of the two ADP·AlF_3_-bound conformations, resembling previously described P1a and P2a states. Subunits are coloured individually. **(B)** Cross-section of the protease showing substrate density (purple) within the axial translocation channel. **(C)** Nucleotide-binding pocket at the subunit interface, showing ADP·AlF_3_ in four catalytic sites and ADP in the remaining sites. **(D)** Substrate density within the AAA^+^ channel engaging the YV-pincer region. The polypeptide (purple) can be modelled in both orientations. Backbone interactions are indicated in yellow.

## Notes

### Competing Interest Statement

The authors have declared no competing interest.

## References

Adams, P.D., Afonine, P.V., Bunkóczi, G., Chen, V.B., Davis, I.W., Echols, N., Headd, J.J., Hung, L.-W., Kapral, G.J., Grosse-Kunstleve, R.W., McCoy, A.J., Moriarty, N.W., Oeffner, R., Read, R.J., Richardson, D.C., Richardson, J.S., Terwilliger, T.C., Zwart, P.H., 2010. PHENIX: a comprehensive Python-based system for macromolecular structure solution. Acta Crystallographica Section D 66, 213–221. 10.1107/S0907444909052925

Bepler, T., Morin, A., Rapp, M., Brasch, J., Shapiro, L., Noble, A.J., Berger, B., 2019. Positive-unlabeled convolutional neural networks for particle picking in cryo-electron micrographs. Nat Methods 16, 1153–1160. 10.1038/s41592-019-0575-8

Biyani, N., Righetto, R.D., McLeod, R., Caujolle-Bert, D., Castano-Diez, D., Goldie, K.N., Stahlberg, H., 2017. Focus: The interface between data collection and data processing in cryo-EM. Journal of Structural Biology 198, 124–133. 10.1016/j.jsb.2017.03.007

Braig, K., Menz, R.I., Montgomery, M.G., Leslie, A.G., Walker, J.E., 2000. Structure of bovine mitochondrial F1-ATPase inhibited by Mg2+ADP and aluminium fluoride. Structure 8, 567–573. 10.1016/S0969-2126(00)00145-3

Chan, D.C., 2020. Mitochondrial Dynamics and Its Involvement in Disease. Annu Rev Pathol 15, 235–259. 10.1146/annurev-pathmechdis-012419-032711

Emsley, P., Cowtan, K., 2004. Coot: model-building tools for molecular graphics. Acta Crystallographica Section D 60, 2126–2132. 10.1107/S0907444904019158

Gibellini, L., Losi, L., De Biasi, S., Nasi, M., Lo Tartaro, D., Pecorini, S., Patergnani, S., Pinton, P., De Gaetano, A., Carnevale, G., Pisciotta, A., Mariani, F., Roncucci, L., Iannone, A., Cossarizza, A., Pinti, M., 2018. LonP1 Differently Modulates Mitochondrial Function and Bioenergetics of Primary Versus Metastatic Colon Cancer Cells. Front. Oncol. 8. 10.3389/fonc.2018.00254

He, L.,Luo Dongyang, Yang,Fan, Li,Chunhao, Zhang,Xuegong, Deng Haiteng, and Zhang, J.-R., 2018. Multiple domains of bacterial and human Lon proteases define substrate selectivity. Emerging Microbes & Infections 7, 1–18. 10.1038/s41426-018-0148-4

Jadiya, P., Tomar, D., 2020. Mitochondrial Protein Quality Control Mechanisms. Genes (Basel) 11, 563. 10.3390/genes11050563

Jr, T.J.H., Trader, D.J., 2025. Exploration of degrons and their ability to mediate targeted protein degradation. RSC Med. Chem. 16, 1067–1082. 10.1039/D4MD00787E

Lee, C., Schwartz, M.P., Prakash, S., Iwakura, M., Matouschek, A., 2001. ATP-dependent proteases degrade their substrates by processively unraveling them from the degradation signal. Mol Cell 7, 627–637. 10.1016/s1097-2765(01)00209-x

López-Otín, C., Blasco, M.A., Partridge, L., Serrano, M., Kroemer, G., 2013. The Hallmarks of Aging. Cell 153, 1194–1217. 10.1016/j.cell.2013.05.039

Lu, B., Lee, J., Nie, X., Li, M., Morozov, Y.I., Venkatesh, S., Bogenhagen, D.F., Temiakov, D., Suzuki, C.K., 2013. Phosphorylation of Human TFAM in Mitochondria Impairs DNA Binding and Promotes Degradation by the AAA+ Lon Protease. Molecular Cell 49, 121–132. 10.1016/j.molcel.2012.10.023

Mohammed, I., Schmitz, K.A., Schenck, N., Balasopoulos, D., Topitsch, A., Maier, T., Abrahams, J.P., 2022. Catalytic cycling of human mitochondrial Lon protease. Structure 30, 1254–1268.e7. 10.1016/j.str.2022.06.006

Ngo, H.B., Kaiser, J.T., Chan, D.C., 2011. The mitochondrial transcription and packaging factor Tfam imposes a U-turn on mitochondrial DNA. Nat Struct Mol Biol 18, 1290–1296. 10.1038/nsmb.2159

Ngo, H.B., Lovely, G.A., Phillips, R., Chan, D.C., 2014. Distinct structural features of TFAM drive mitochondrial DNA packaging versus transcriptional activation. Nat Commun 5, 3077. 10.1038/ncomms4077

Ngo, J.K., Davies, K.J.A., 2009. Mitochondrial Lon protease is a human stress protein. Free Radical Biology and Medicine 46, 1042–1048. 10.1016/j.freeradbiomed.2008.12.024

Okuno, T., Yamanaka, K., Ogura, T., 2006. An AAA protease FtsH can initiate proteolysis from internal sites of a model substrate, apo-flavodoxin. Genes to Cells 11, 261–268. 10.1111/j.1365-2443.2006.00940.x

Ondrovičová, G., Liu, T., Singh, K., Tian, B., Li, H., Gakh, O., Perečko, D., Janata, J., Granot, Z., Orly, J., Kutejová, E., Suzuki, C.K., 2005. Cleavage Site Selection within a Folded Substrate by the ATP-dependent Lon Protease*. Journal of Biological Chemistry 280, 25103–25110. 10.1074/jbc.M502796200

Parsell, D.A., Sauer, R.T., 1989. The structural stability of a protein is an important determinant of its proteolytic susceptibility in Escherichia coli. Journal of Biological Chemistry 264, 7590–7595. 10.1016/S0021-9258(18)83275-6

Punjani, A., Rubinstein, J.L., Fleet, D.J., Brubaker, M.A., 2017. cryoSPARC: algorithms for rapid unsupervised cryo-EM structure determination. Nat Methods 14, 290–296. 10.1038/nmeth.4169

Rigas, S., Daras Gerasimos, Sweetlove,Lee J., and Hatzopoulos, P., 2009. Mitochondria biogenesis via Lon1 selective proteolysis: Who dares to live for ever? Plant Signaling & Behavior 4, 221–224. 10.4161/psb.4.3.7863

Shin, M., Watson, E.R., Song, A.S., Mindrebo, J.T., Novick, S.J., Griffin, P.R., Wiseman, R.L., Lander, G.C., 2021. Structures of the human LONP1 protease reveal regulatory steps involved in protease activation. Nat Commun 12, 3239. 10.1038/s41467-021-23495-0

Strauss, K.A., Jinks, R.N., Puffenberger, E.G., Venkatesh, S., Singh, K., Cheng, I., Mikita, N., Thilagavathi, J., Lee, J., Sarafianos, S., Benkert, A., Koehler, A., Zhu, A., Trovillion, V., McGlincy, M., Morlet, T., Deardorff, M., Innes, A.M., Prasad, C., Chudley, A.E., Lee, I.N.W., Suzuki, C.K., 2015. CODAS Syndrome Is Associated with Mutations of LONP1, Encoding Mitochondrial AAA+ Lon Protease. The American Journal of Human Genetics 96, 121–135. 10.1016/j.ajhg.2014.12.003

Szczepanowska, K., Trifunovic, A., 2022. Mitochondrial matrix proteases: quality control and beyond. The FEBS Journal 289, 7128–7146. 10.1111/febs.15964

Tang, G., Peng, L., Baldwin, P.R., Mann, D.S., Jiang, W., Rees, I., Ludtke, S.J., 2007. EMAN2: An extensible image processing suite for electron microscopy. Journal of Structural Biology, Software tools for macromolecular microscopy 157, 38–46. 10.1016/j.jsb.2006.05.009

Taouktsi, E., Kyriakou, E., Smyrniotis, S., Borbolis, F., Bondi, L., Avgeris, S., Trigazis, E., Rigas, S., Voutsinas, G.E., Syntichaki, P., 2022. Organismal and Cellular Stress Responses upon Disruption of Mitochondrial Lonp1 Protease. Cells 11, 1363. 10.3390/cells11081363

Varshavsky, A., 2011. The N-end rule pathway and regulation by proteolysis. Protein Sci 20, 1298–1345. 10.1002/pro.666

Venkatesh, S., Lee, J., Singh, K., Lee, I., Suzuki, C.K., 2012. Multitasking in the mitochondrion by the ATP-dependent Lon protease. Biochim Biophys Acta 1823, 56–66. 10.1016/j.bbamcr.2011.11.003

Xu, Y.-W., Moréra, S., Janin, J., Cherfils, J., 1997. AlF3 mimics the transition state of protein phosphorylation in the crystal structure of nucleoside diphosphate kinase1and1MgADP. Proc Natl Acad Sci U S A 94, 3579–3583. 10.1073/pnas.94.8.3579

Yang, Z., Fang, J., Chittuluru, J., Asturias, F.J., Penczek, P.A., 2012. Iterative Stable Alignment and Clustering of 2D Transmission Electron Microscope Images. Structure 20, 237–247. 10.1016/j.str.2011.12.007

